## Supplemental Figures for "Directed differentiation of mouse pluripotent stem cells into functional lung-specific mesenchyme"

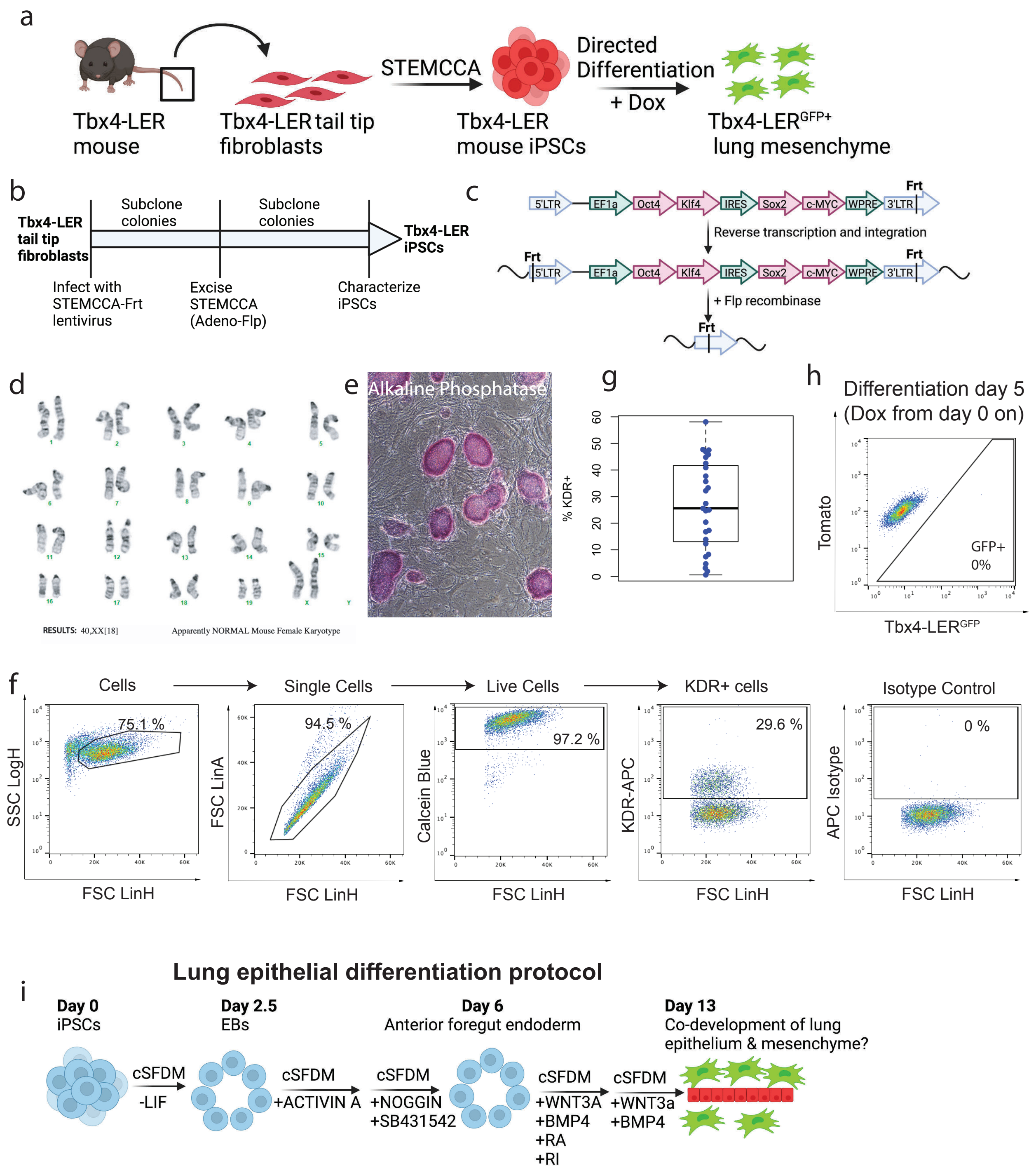


**Supplementary Figure 1: Reprograming of Tbx4-LER$^{GFP}$ tail tip fibroblasts and directed lung mesenchymal differentiation.**

**a:** Overview of reprogramming procedure and generation of induced lung mesenchyme.

**b:** Schematic showing the generation of iPSCs from tail tip fibroblasts.

**c:** Schematic showing STEMCCA lentiviral cassette before and after excision of the Frt-flanked STEMCCA sequence.

**d:** Karyotyping results showing normal karyotype of Tbx4-LER iPSC line.

**e:** Alkaline phosphatase stain of Tbx4-LER iPSC line.

**f:** Example of gating strategy and representative flow cytometry plot showing KDR stain and isotype on day 5 of differentiation.

**g:** Percentage of KDR+ cells on day 5 of differentiation (lateral plate mesoderm stage). Each dot represents an individual differentiation experiment.

**h:** Flow cytometry plot showing expression of GFP and Tomato on day 5 of differentiation, when dox was kept in the medium from day 0 on.

**i:** Protocol for the directed differentiation of iPSCs towards the lung epithelial lineage. cSFDM = complete serum-free differentiation medium, RA = retinoic acid, RI = rock inhibitor (Y-27632).

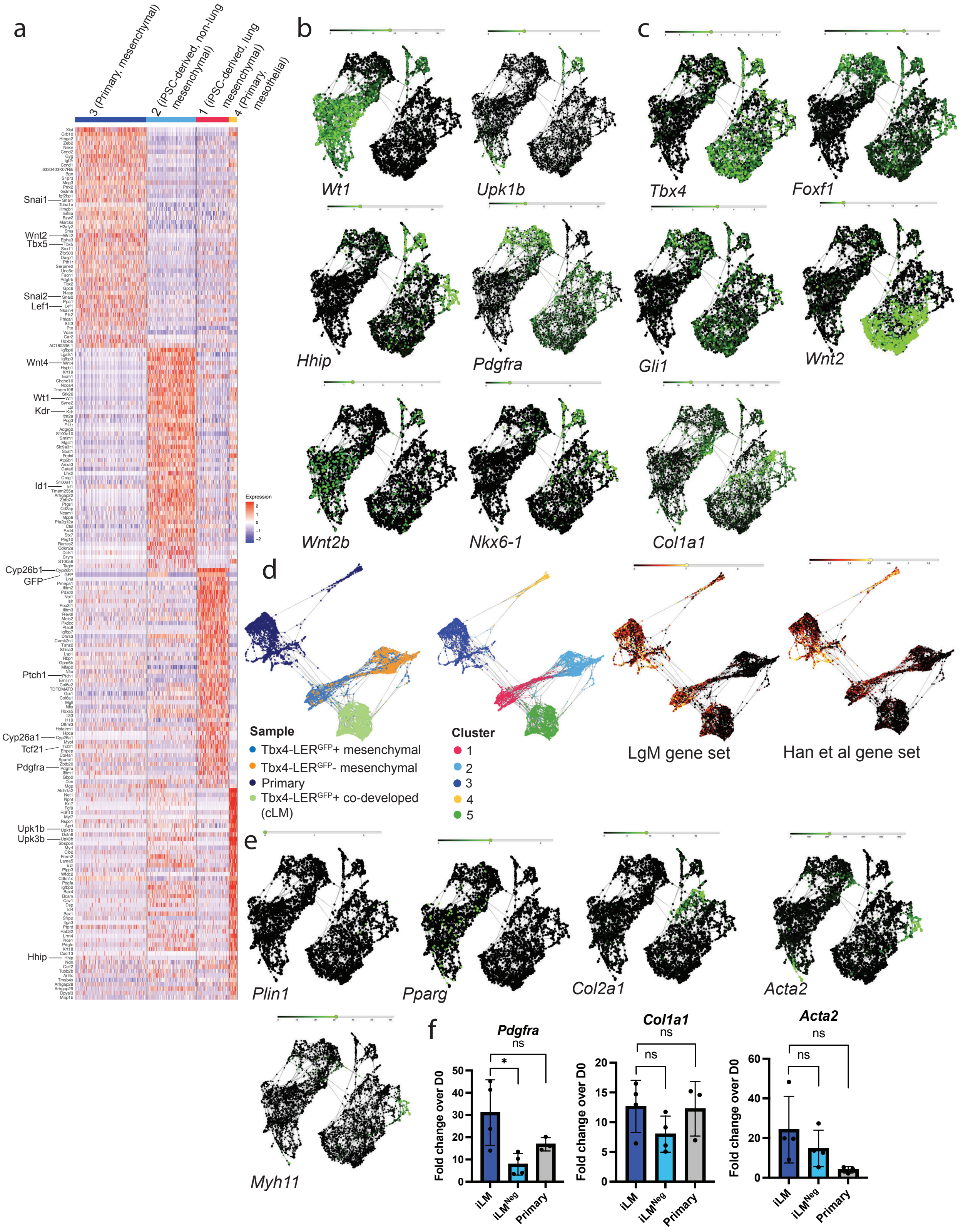


**Supplementary Figure 2: Single cell RNA sequencing of iPSC-derived and primary lung mesenchyme.**

**a:** Heatmap showing top 50 enriched genes in clusters previously defined by Louvain clustering (resolution 0.1). Genes of interest are highlighted in larger font.
**b:** SPRING plot showing expression of mesothelial markers Wt1 and Upk1b in iPSC-derived and primary cell clusters.
**c:** SPRING plot showing expression of lung mesenchymal markers of interest in iPSC-derived and primary cell clusters.
**d:** SPRING plot overview, Louvain clusters (resolution 0.1), and enrichment of the LgM and Han et al. gene sets in scRNA-seq dataset including Tbx4-LER$^{GFP}$+/- generated with the lung mesenchymal differentiation protocol ("mesenchymal"), as well as Tbx4-LER$^{GFP}$+ cells generated by co-development ("co-developed").
**e:** SPRING plot showing expression of mature mesenchymal lineage markers in iPSC-derived and primary cell clusters (*Plin*, *Pparg* = adipogenic/lipofibroblast; *Col2a1* = chondrogenic; *Acta2*, *Myh11* = smooth muscle).
**f:** RT-qPCR showing fold change expression relative to day 0 iPSCs of additional lung mesenchyme markers in induced lung mesenchyme (iLM) and iLM$^{Neg}$ cells compared to primary lung mesenchyme from E12.5 embryos. Tbx4-LER$^{GFP}$+/- cells were collected on differentiation day 13 and dox was added from day 5 on. N ≥ 3. Bars show mean ± sd. *$p<0.05$, **$p<0.01$, ***$p<0.001$, ns = non-significant, as determined by unpaired, two-sided Student's t test.

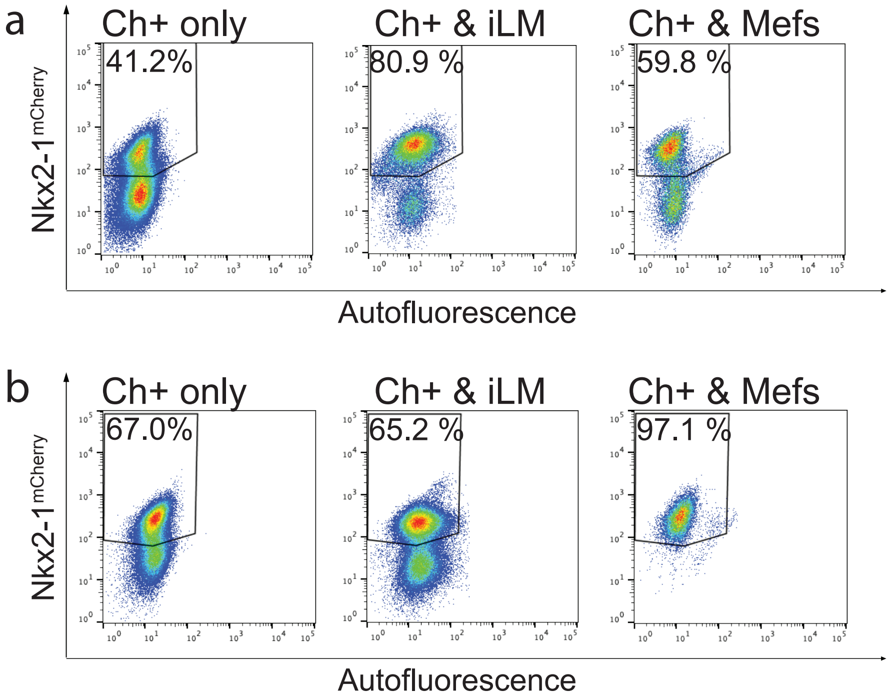


**Supplementary Figure 3: Flow cytometry analysis of co-cultures of induced lung mesenchyme with Nkx2-1$^{mCherry}$+ lung epithelial progenitors in distal and proximal epithelial medium.**

**a:** Representative flow cytometry plots showing percentage of Nkx2-1$^{mCherry}$+ cells for Nkx2-1$^{mCherry}$+ cells cultured alone, with iLM or with MEFs for 7 days in distal medium. Cells were pre-gated for single cells, live cells, GFP- and Epcam+ cells.

**b:** Representative flow cytometry plots showing percentage of Nkx2-1$^{mCherry}$+ cells for Nkx2-1$^{mCherry}$+ cells cultured alone, with iLM or with MEFs for 7 days in proximal medium. Cells were pre-gated for single cells, live cells, GFP- and Epcam+ cells.

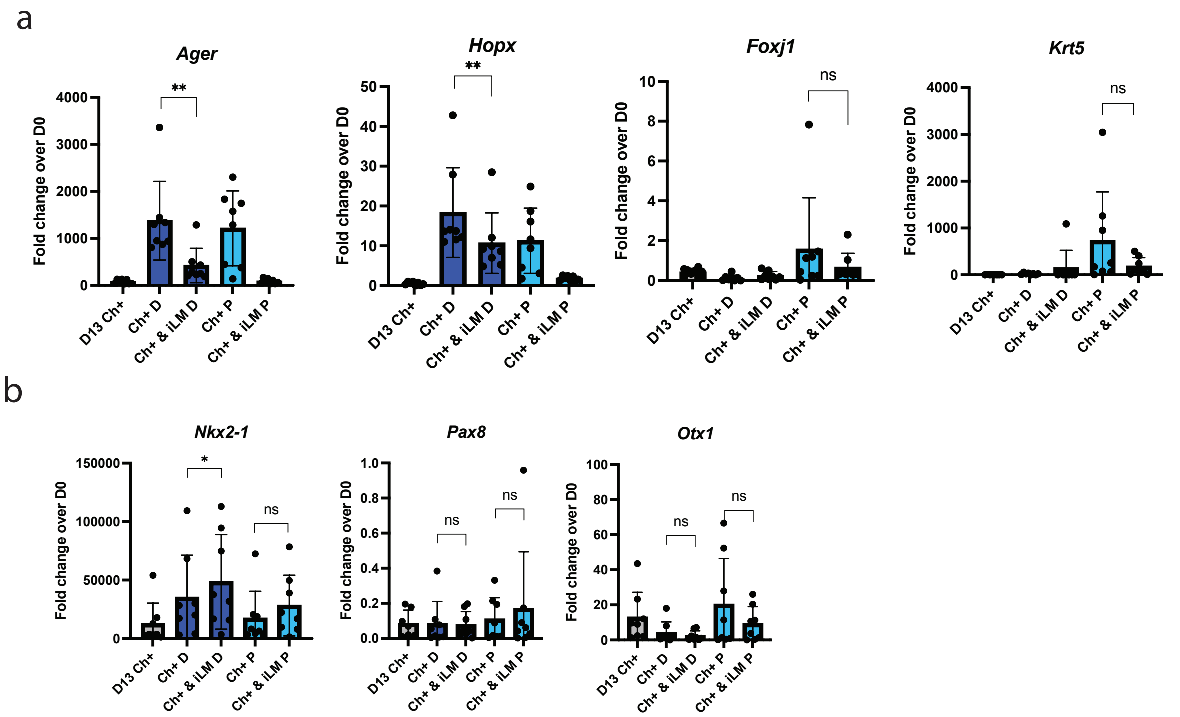


**Supplementary Figure 4: Gene expression analysis of co-cultured lung epithelial progenitors.**
**a:** RT-qPCR data showing fold change expression relative to day 0 iPSCs of additional distal and proximal lung/airway markers in Nkx2-1$^{mCherry}$+ cells before (grey) and after co-culture in distal (dark blue) and proximal (light blue) medium. N = 8.
**b:** RT-qPCR data showing fold change expression relative to day 0 iPSCs of *Nkx2-1*, *Pax8*, and *Otx1* in Nkx2-1$^{mCherry}$+ cells before (grey) and after co-culture in distal (dark blue) and proximal (light blue) medium. N = 8.
All bars show mean ± sd. *p<0.05, **p<0.01, ***p<0.001, ns = non-significant, as determined by paired, two-sided Student's t test.

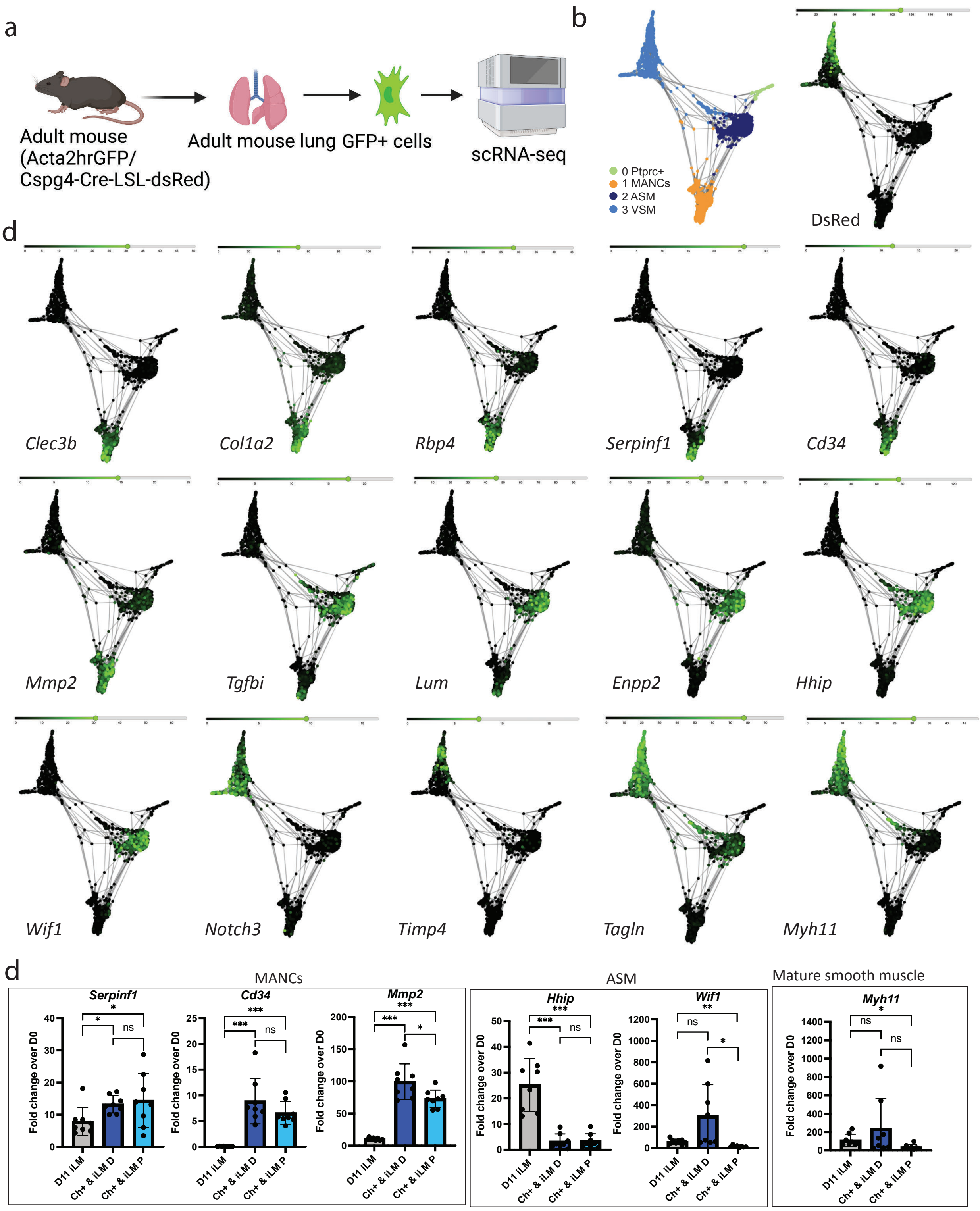


**Supplementary Figure 5: scRNA-seq of adult Acta2$^{GFP}$+ cells and gene expression analysis of co-cultured iLM.**

**a:** Schematic of scRNAseq of Acta2$^{GFP}$+ cells from adult mouse lungs.
**b:** SPRING plots showing cluster annotation and expression of DsRed in scRNA-seq dataset.
**c:** SPRING plots showing expression of MANC, ASM and VSM markers used for RT-qPCR in scRNA-seq dataset.
**d:** RT-qPCR data showing fold change expression relative to day 0 iPSCs of additional MANC, ASM and mature smooth muscle markers in day 11 iLM cells before co-culture (grey), and in iLM after co-culture in distal (dark blue) and proximal (light blue) medium. N = 8. All bars show mean ± sd. *$p<0.05$, **$p<0.01$, ***$p<0.001$, ns = non-significant, as determined by paired, two-sided Student's t test.

**Supplementary Table 1: Embryonic lung mesenchymal gene sets.**
Gene set lists used to test for enrichment of lung mesenchymal genes. LgM gene set consists of genes previously suggested to be expressed in early lung mesenchyme, Han et al. gene set consist of a list of genes found to be enriched in E9.5 mouse lung mesenchyme by Han et al, 2020.

| **LgM gene set** | **Han et al. gene set** |
|---|---|
| *Wnt2* | *Osr1* |
| *Wnt2b* | *Nkx6-1* |
| *Gli1* | *Rspo1* |
| *Gli2* | *Smpx* |
| *Gli3* | *Stmn1* |
| *Hhip* | *Tbx5* |
| *Ptch1* | *Cxcl13* |
| *Foxf1* | *Nr2f2* |
| *Tbx4* | *Hoxa5* |
| *Tbx5* | *Srsf2* |
| *Fgf10* | *Msc* |
| *Pdgfra* | *Sparcl1* |
| *Nkx6-1* | *Foxf1* |
| | *Pde5a* |
| | *Arl4a* |
| | *Hmga2* |
| | *Foxp1* |
| | *Nfib* |
| | *Ptms* |
| | *Dbi* |
| | *Fendrr* |
| | *Tnfrsf19* |
| | *Npc2* |
| | *Zfp503* |
| | *Fat1* |
| | *Id3* |
| | *Bdnf* |
| | *Vgll4* |
| | *Chd3* |
| | *Hoxa4* |
| | *Dpysl3* |
| | *Bcl11a* |
| | *Nrep* |
| | *Mllt3* |
| | *Serf1* |
| | *Unc5c* |
| | *Manf* |

| | |
|---|---|
| | *Nkd1* |
| | *Wnt2* |
| | *Nrip1* |
| | *Tbx3* |
| | *Mt1* |
| | *Foxp2* |
| | *Lxn* |
| | *Cdk6* |
| | *Tanc1* |
| | *Pdgfc* |
| | *Pitx2* |
| | *Igfbp2* |
| | *Auts2* |
| | *Fras1* |
| | *Mfge8* |
| | *Cacnb3* |
| | *Pgf* |
| | *Uchl1* |
| | *Sfrp2* |
| | *Adgrl2* |
| | *Xbp1* |
| | *Apoe* |
| | *Xist* |
| | *Gng2* |

**Supplementary Table 2: Top 25 enriched genes in MANC, ASM, and VSM cells in Zepp et al 2021.** Top genes enriched genes used for gene sets in Figure 7 D.

| MANCs | ASM | VSM |
|---|---|---|
| *Serpinf1* | *Hhip* | *Actg2* |
| *Cygb* | *Lum* | *Lmod1* |
| *Pi16* | *Igfbp5* | *Adam33* |
| *Rbp4* | *Enpp2* | *Acta2* |
| *Smoc2* | *Fgf18* | *Myh11* |
| *Dcn* | *Aspn* | *Map3k7cl* |
| *Entpd2* | *Tgfbi* | *Cnn1* |
| *Tspan11* | *Cdh4* | *Fam129a* |
| *Igfbp4* | *Mustn1* | *Tpm2* |
| *Col14a1* | *6330403K07Rik* | *Slc11a1* |
| *Scara5* | *Pde10a* | *Map1b* |
| *Fbln1* | *Gja1* | *Col8a1* |
| *Pdgfrl* | *P2ry14* | *Actc1* |
| *Mmp23* | *P4ha3* | *Aoc3* |
| *Col1a2* | *Cntfr* | *Ramp1* |
| *Col15a1* | *Cyp2e1* | *Mustn1* |
| *Clec3b* | *Grem2* | *Myl9* |
| *F3* | *Sgcg* | *Cdh13* |
| *Ccl11* | *Cd9* | *Sh3bgr* |
| *Itgbl1* | *Wif1* | *Gpc6* |
| *Igfbp6* | *Fxyd6* | *Aspn* |
| *Col1a1* | *Ngb* | *Tagln* |
| *Adamts2* | *Clu* | *Crip1* |
| *Fgl2* | *Ptma* | *Des* |

**Supplementary Table 3: Enriched genes in Acta2+ clusters.**
Top 30 genes enriched in MANC, ASM and VSM cells in our scRNA-seq dataset.

| MANCs | ASM | VSM |
|---|---|---|
| *Gsn* | *Hhip* | *Myl6* |
| *Igfbp6* | *Lum* | *Myl9* |
| *Serping1* | *Bmp5* | *Tpm2* |
| *Mfap4* | *Enpp2* | *Acta2* |
| *Col1a2* | *Malat1* | *Myh11* |
| *Inmt* | *Gja1* | *Tagln* |
| *Clec3b* | *Thbs1* | *Ppp1r14a* |
| *Mmp2* | *Wif1* | *Tpm1* |
| *Cfh* | *Rarres2* | *Dstn* |
| *Cpxm1* | *Aspn* | *Des* |
| *Rbp1* | *Dpt* | *Flna* |
| *Olfml3* | *Mt1* | *Actg2* |
| *Dpep1* | *Igfbp5* | *Tinagl1* |
| *Cd34* | *P4ha3* | *Cald1* |
| *Pid1* | *Mt2* | *Epas1* |
| *Pcolce* | *Odc1* | *Mylk* |
| *Nbl1* | *Tgfbi* | *Pbxip1* |
| *Efemp1* | *Pcsk5* | *Mcam* |
| *Abca8a* | *Pappa2* | *Actn1* |
| *Cygb* | *Pam* | *Tns1* |
| *Adh1* | *Cdh4* | *Sncg* |
| *Gpc3* | *S100b* | *Mustn1* |
| *Timp2* | *Cd44* | *Notch3* |
| *Itgbl1* | *Sema3c* | *Itgb1* |
| *C3* | *6330403K07Rik* | *Fam129a* |
| *Rbp4* | *Adamtsl2* | *Ppp1r12a* |
| *Adamts2* | *Sgcg* | *Lpp* |
| *Slc43a3* | *Fmo2* | *Lmod1* |
| *Rnase4* | *Rnf122* | *Rbpms* |
| *Serpinf1* | *Cntfr* | *Map1b* |
